## Supplemental information for "Single-nucleus multiomics reveals the gene-regulatory networks underlying sex determination of murine primordial germ cells"

**Supplementary Figure 1**

**Supplementary Figure 2**

**Supplementary Figure 3**

**Supplementary Figure 4**

**Supplementary Figure 5**

**Supplementary Figure 6**

**Supplementary Figure 7**

**Supplementary Figure 8**

**Supplementary Figure 9**

**Supplementary Figure 10**

**Supplementary Figure 11**

**Supplementary Figure 12**

### Supplementary Figure 1

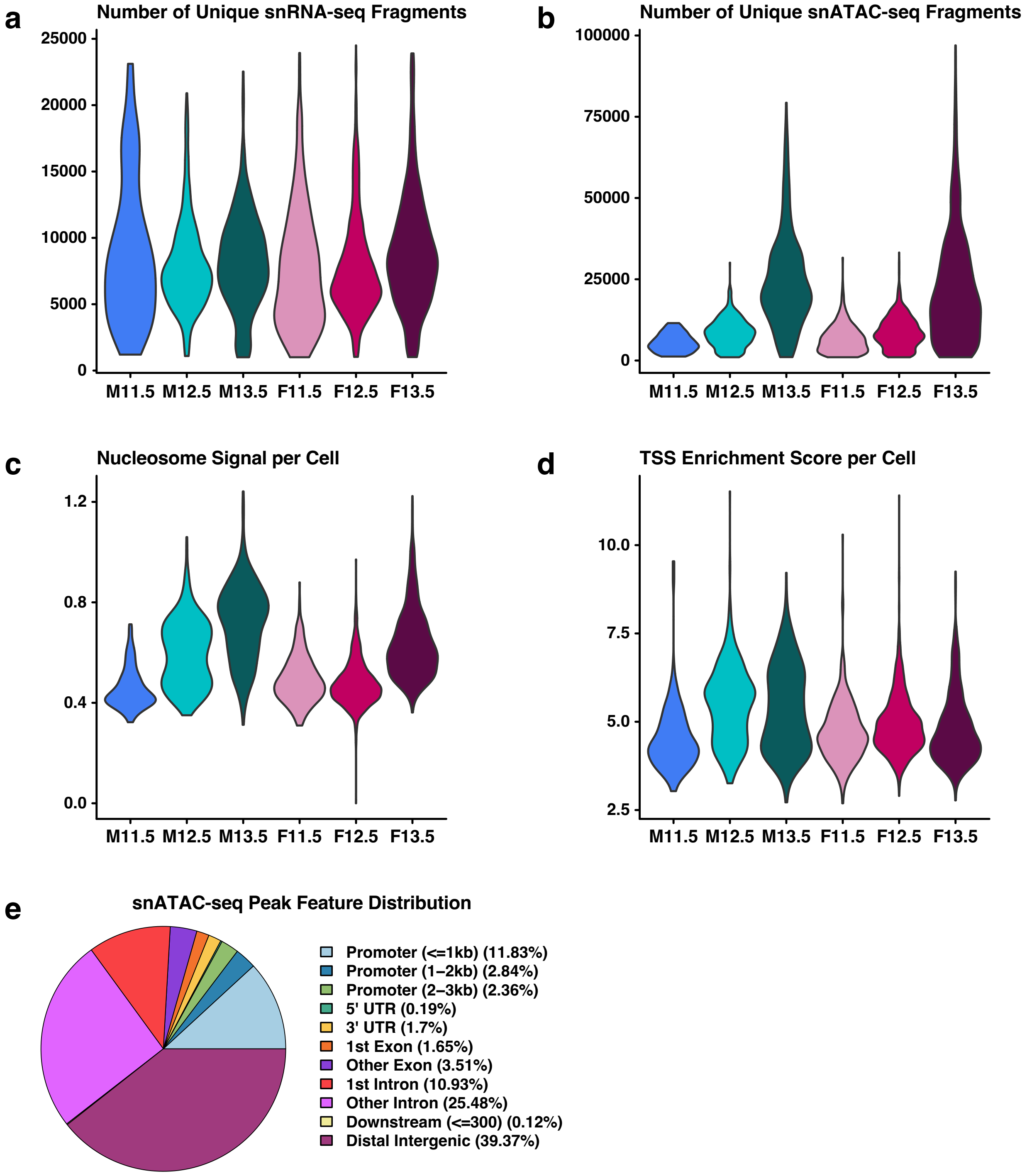


**Supplementary Figure 1: QC statistics for the snRNA-seq and snATAC-seq libraries of E11.5-E13.5 XX and XY PGCs.** (a) Violin plot of the number of unique snRNA-seq reads per cell for E11.5-E13.5 XX and XY PGCs. (b) Violin plot of the number of unique snATAC-seq fragments per cell for E11.5-E13.5 XX and XY PGCs. (c) snATAC-seq nucleosome signal per cell for E11.5-E13.5 XX and XY PGCs. (d) snATAC-seq transcription start site (TSS) enrichment score per cell for E11.5-E13.5 XX and XY PGCs. (e) Feature distribution of snATAC-seq peaks called for E11.5-E13.5 XX and XY PGCs.

**Supplementary Figure 2**

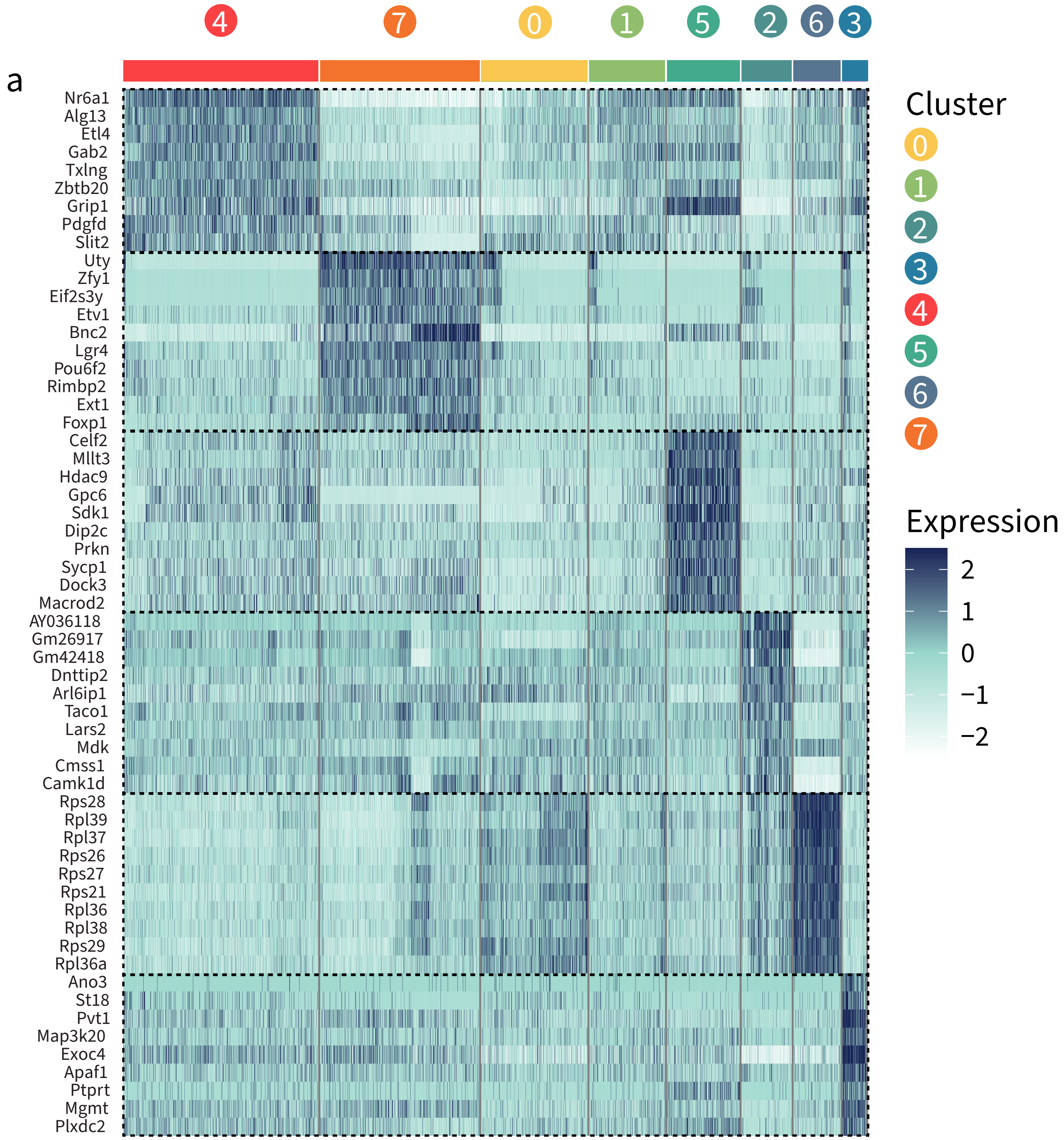


**Supplementary Figure 2: Heatmap of marker gene expression for snRNA-seq clusters of E11.5-E13.5 XX and XY PGCs.** (a) Heatmap of marker gene expression for unbiasedly identified snRNA-clusters of E11.5-E13.5 XX and XY PGCs. The color scale represents the expression level of each gene. The colored bar at top of heatmap represents the cluster number.

**Supplementary Figure 3**

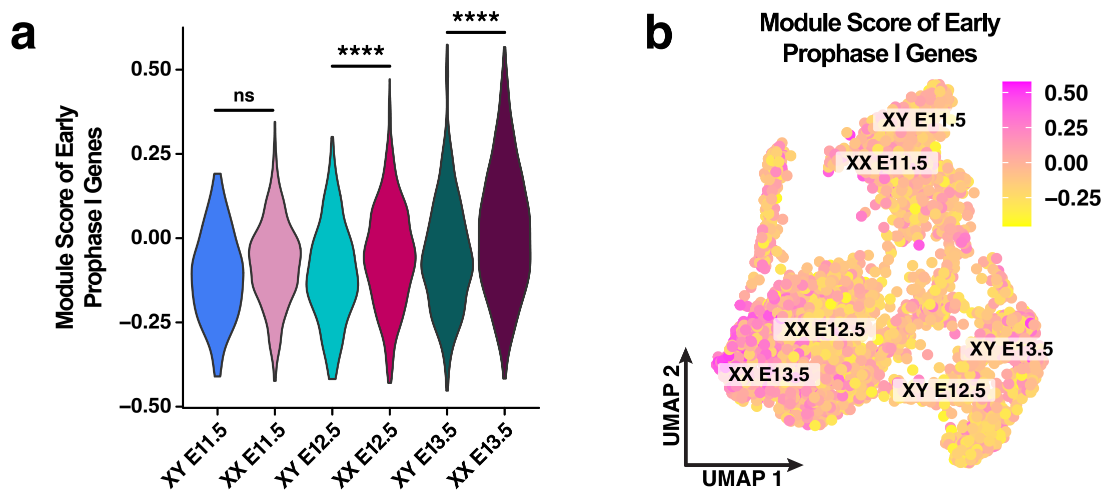


**Supplementary Figure 3: Module score of early prophase I genes.** (a) Violin plot of the module score, or average expression, of early prophase I genes for each PGC population clustered by embryonic stage and sex. ****$p < 0.0001$; One-Way ANOVA with post-hoc Tukey multiple comparison test. (b) Joint UMAP of integrated snRNA-seq and snATAC-seq data showing the module score levels of early prophase I genes across all PGC populations. (a-b) Early prophase I genes included *Rad21, Rad21l, Rec8, Ccnb3, Sycp2, Rad51, Hormad, Sycp1,* and *Kit*.

### Supplementary Figure 4

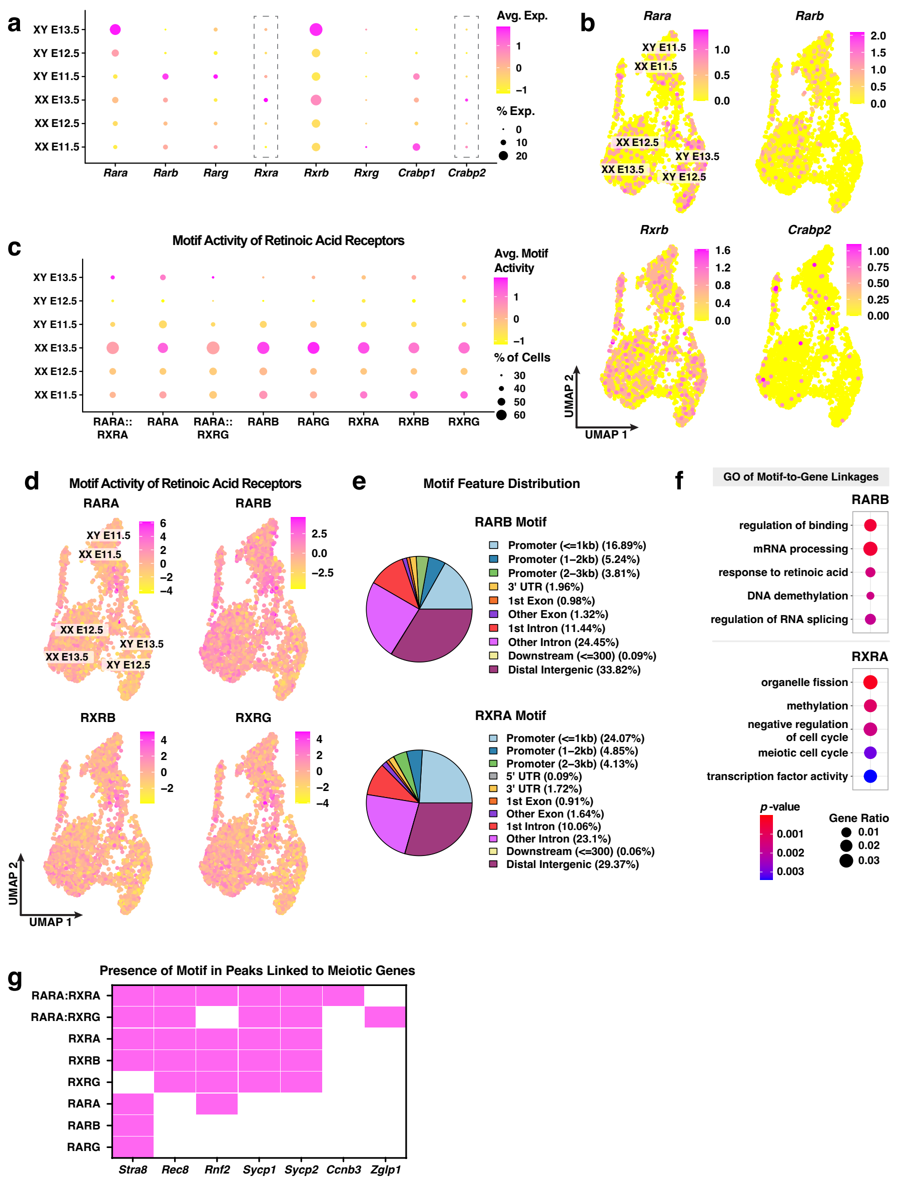


**Supplementary Figure 4: Regulatory potential of retinoic acid receptors.** (a) Dotplot of the average expression and percentage of cells expressing *Rara, Rarb, Rarg, Rxra, Rxrb, Rxrg, Crabp1,* and *Crabp2*. The color scale represents the average expression level and the size of the dot represents the percentage of cells expressing the gene. (b) Joint UMAP showing *Rara, Rarb, Rxrb,* and *Crabp2* expression levels for E11.5-E13.5 XX and XY PGCs. Individual cells are color coded by the level of gene expression. (c) *In silico* chromatin binding score, termed 'motif activity', of retinoic acid receptors. Average motif activity is indicated by the color scale and the percentage of cells is indicated by the size of the dot. (d) Joint UMAP showing the motif activity scores of RARA, RARB, RXRB, and RXRG for E11.5-E13.5 XX and XY PGCs. (e) Pie chart showing the feature distribution of the RARB and RXRA motifs. (f) Gene ontology (GO) of predicted target genes of RARB and RXRA based on motif-to-gene linkages. The color scale represents the *p*-value of GO term enrichment, and the size of the dot indicates the gene ratio for each GO term. (g) Icon array showing the presence (pink) or absence (white) of the RA receptor motif (*y*-axis) in peaks linked to meiotic genes.

### Supplementary Figure 5

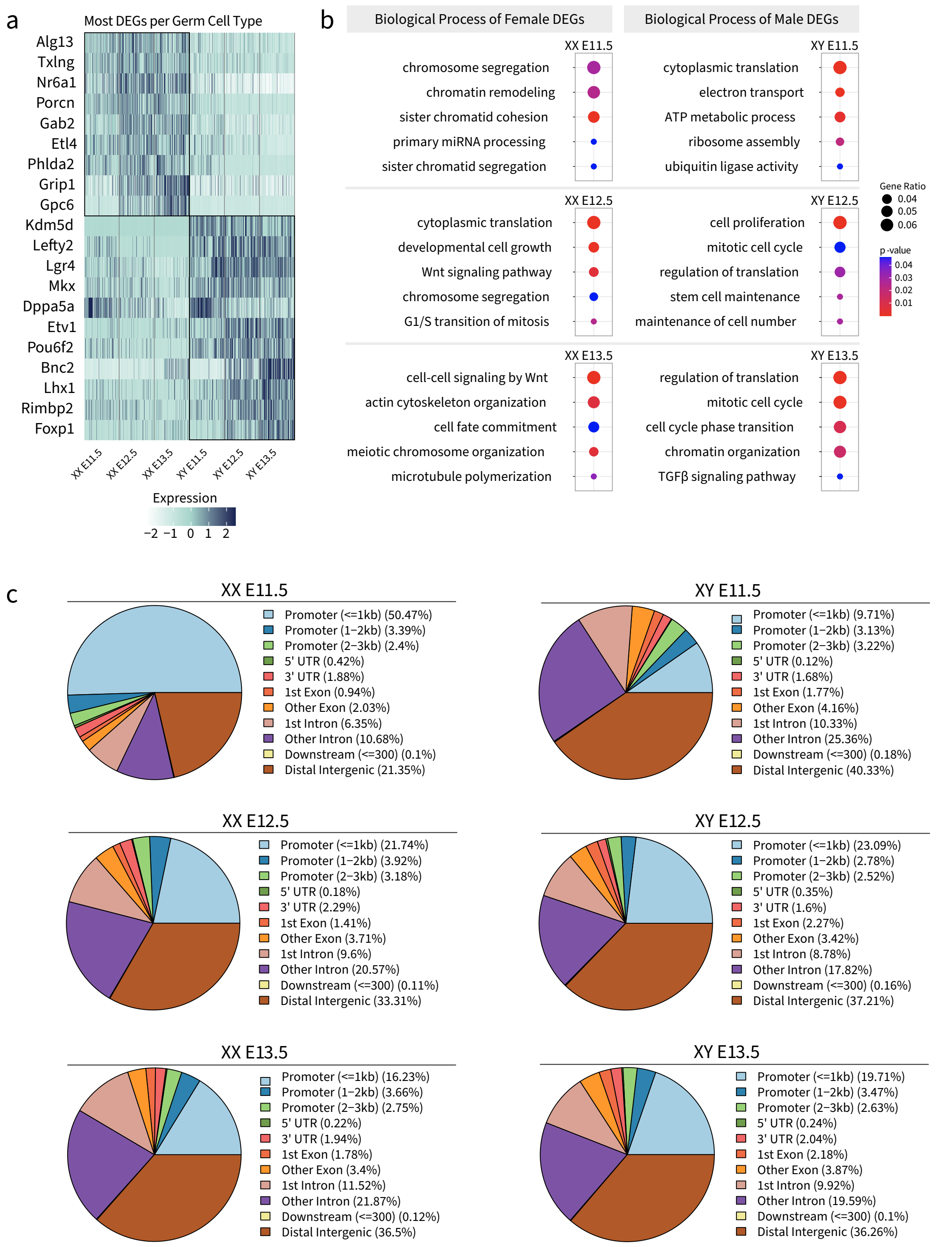


**Supplementary Figure 5: Characterization of differentially expressed genes and differentially accessible peaks in E11.5-E13.5 XX and XY PGCs.** (a) Heatmap of the most differentially expressed genes (DEGs) for each PGC population. The color scale represents the level of expression. (b) Biological process gene ontology (GO) analysis of XX and XY DEGs. The color scale represents the *p*-value of GO term enrichment, and the size of the dot indicates the gene ratio for each GO term. (c) Pie charts showing the feature distribution of differentially accessible chromatin peaks for E11.5-E13.5 XX and XY PGCs.

### Supplementary Figure 6

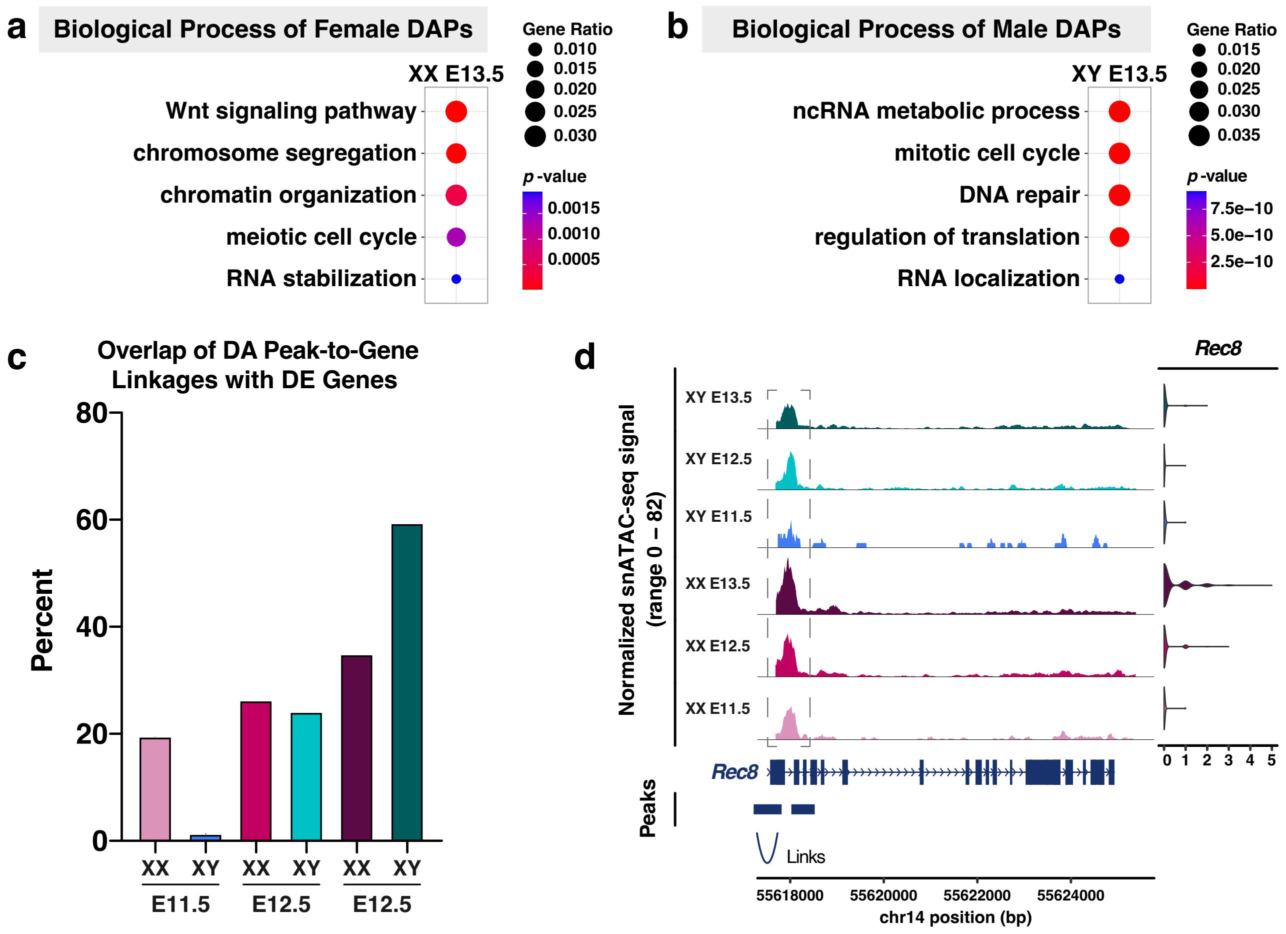


**Supplementary Figure 6: Characterization of peak-to-gene linkages in E11.5-E13.5 XX and XY PGCs.** (a-b) Biological process gene ontology (GO) analysis of E13.5 XX (a) and XY (b) differentially accessible peaks (DAPs). The color scale represents the *p*-value of GO term enrichment, and the size of the dot indicates the gene ratio for each GO term. (c) Bar plot showing the percentage of DAPs with peak-to-gene linkages to differentially expressed genes. (d) *Left:* Coverage plot of the normalized snATAC-seq signal at the *Rec8* locus. Peak-to-gene linkages are indicated by the 'links' line. *Right:* Violin plot of expression in E11.5-E13.5 XX and XY PGCs.

### Supplementary Figure 7

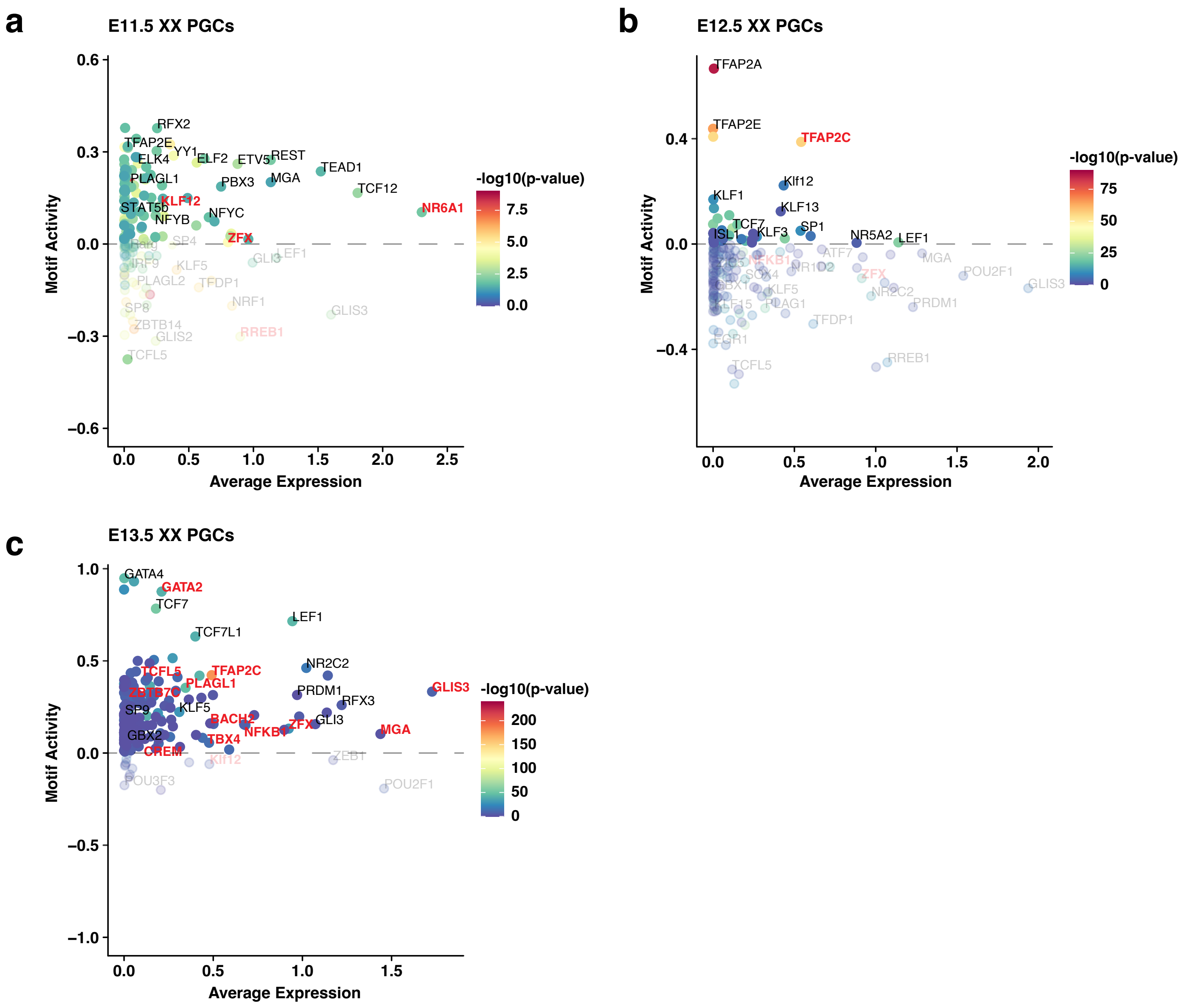


**Supplementary Figure 7: Identification of enriched transcription factors in XX PGCs.** (a-c) *In silico* chromatin binding score, termed 'motif activity', vs average expression of transcription factors (TFs) that bind significantly enriched motifs in E11.5 (a), E12.5 (b), and E13.5 (c) XX PGCs. The color scale represents the -$\log_{10}$($p$-value) of motif enrichment. TF names colored in red are significantly upregulated in XX PGCs when compared to XY PGCs.

### Supplementary Figure 8

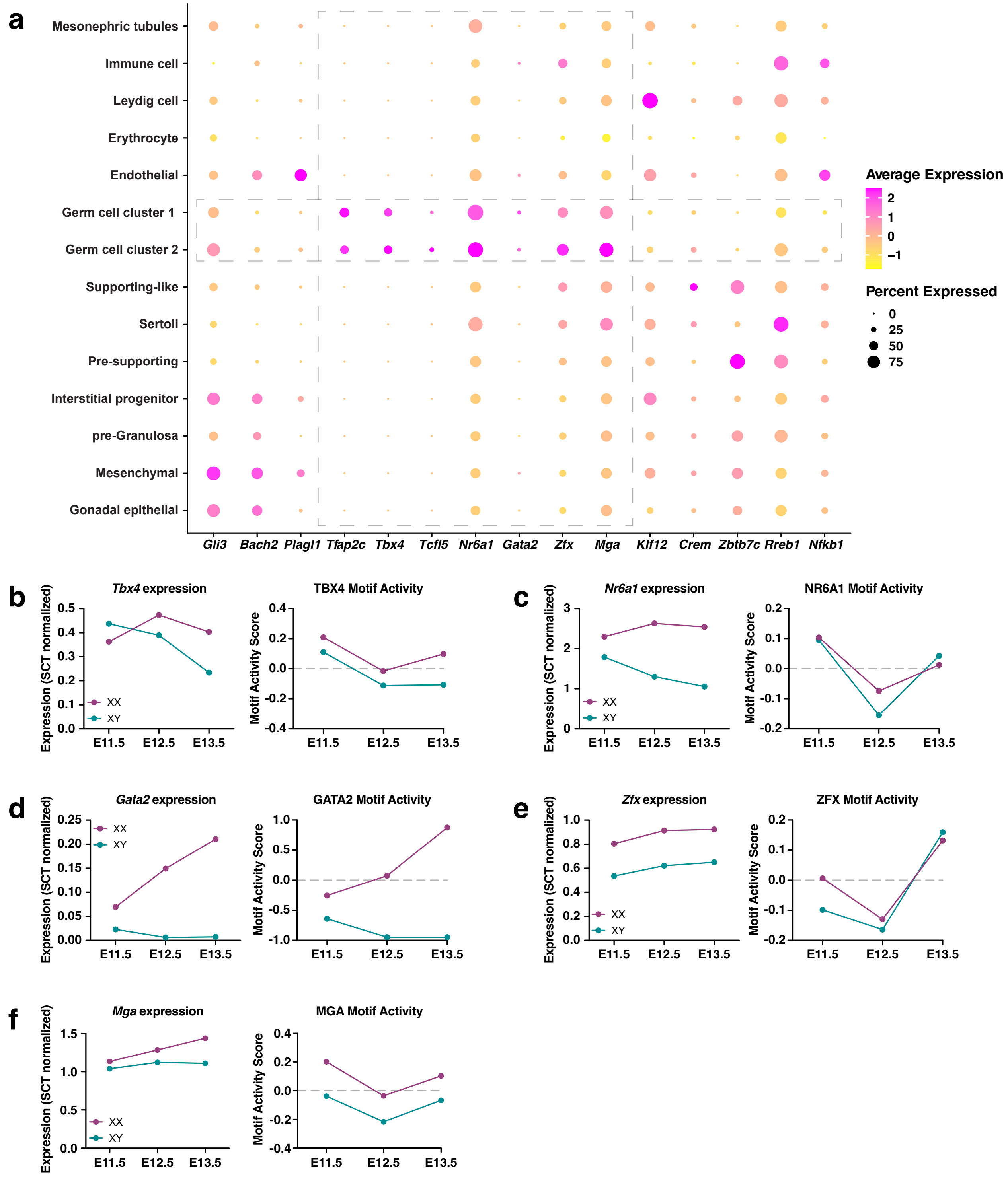


**Supplementary Figure 8: Identification of strong transcription factor candidates in XX PGCs.** (a) Dotplot of the average expression and percentage of cells expressing XX PGC-enriched transcription factors (TFs) in PGCs and gonadal somatic cells. Dashed boxes indicate TFs that are enriched in PGCs when compared to the somatic compartment. The color scale represents the average expression level, and the size of the dot represents the percentage of cells expressing the gene. (b-f) Line plots showing the expression levels and *in silico* chromatin binding, termed 'motif activity', of *Tbx4* (b), *Nr6a1* (c), *Gata2* (d), *Zfx* (d), and *Mga* (f) in E11.5-E13.5 XX (pink) and XY (teal) PGCs.

### Supplementary Figure 9

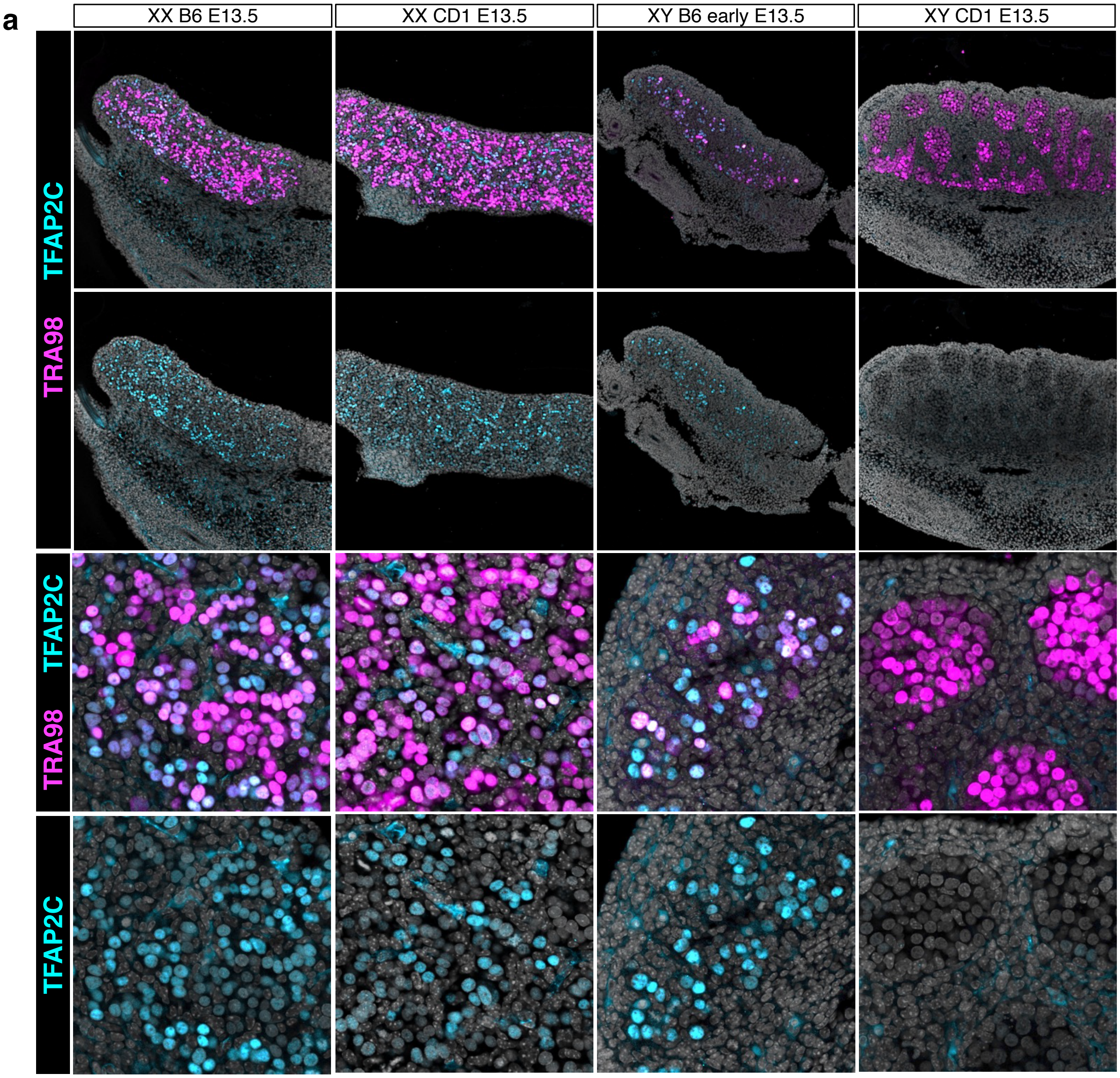


**Supplementary Figure 9: Immunofluorescence staining of TFAP2C in E13.5 gonads.** (a) DAPI staining (grey) and immunofluorescence staining of TRA98 (pink) and TFAP2C (blue) in E13.5 gonads from XX and XY mice on the C57Bl/6 or CD1 background.

### Supplementary Figure 10

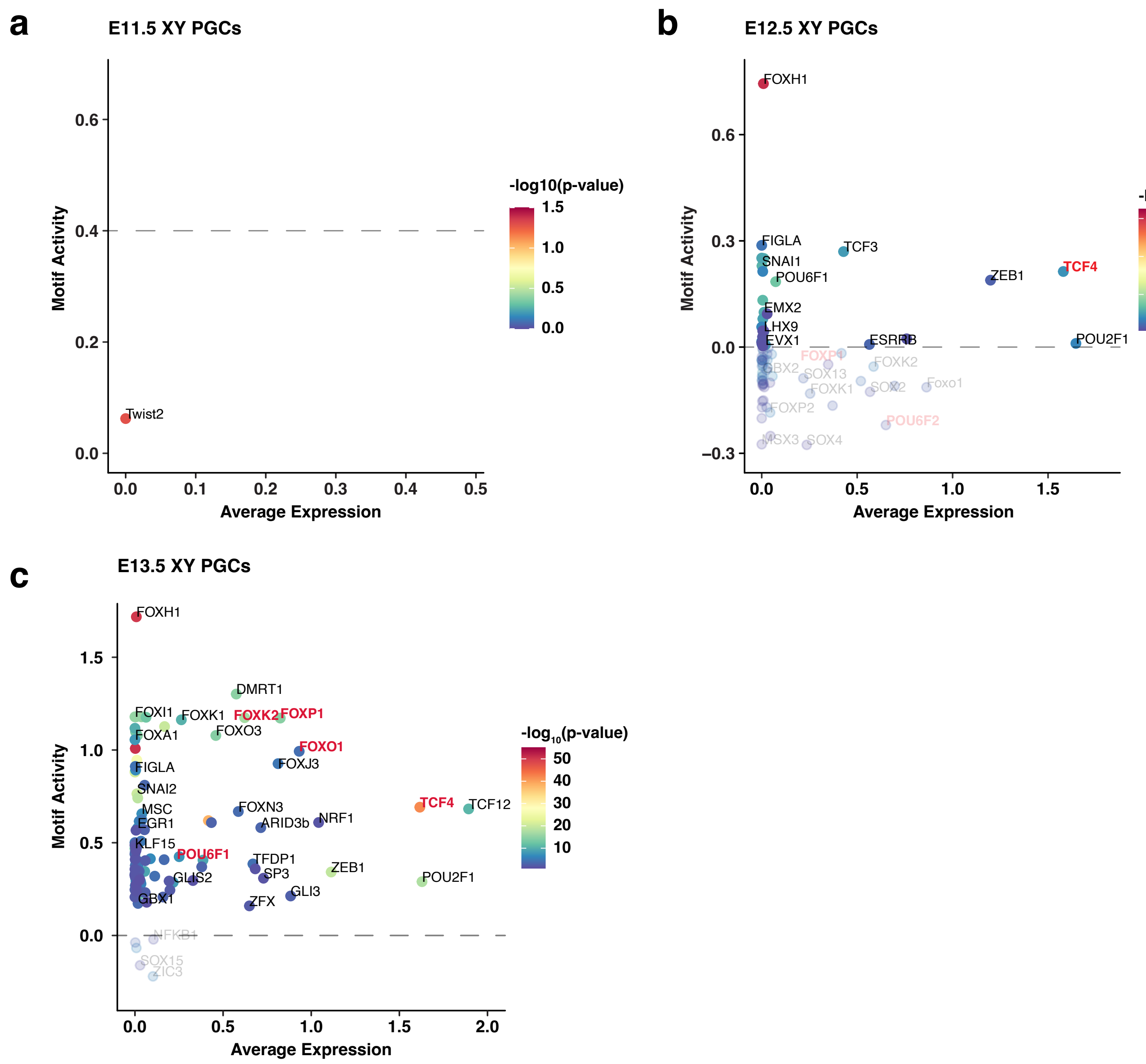


**Supplementary Figure 10: Identification of enriched transcription factors in XY PGCs.** (a-c) *In silico* chromatin binding score, termed 'motif activity', vs average expression of transcription factors (TFs) that bind significantly enriched motifs in E11.5 (a), E12.5 (b), and E13.5 (c) XY PGCs. The color scale represents the -$\log_{10}$($p$-value) of motif enrichment. TF names colored in red are significantly upregulated in XY PGCs when compared to XX PGCs.

### Supplementary Figure 11

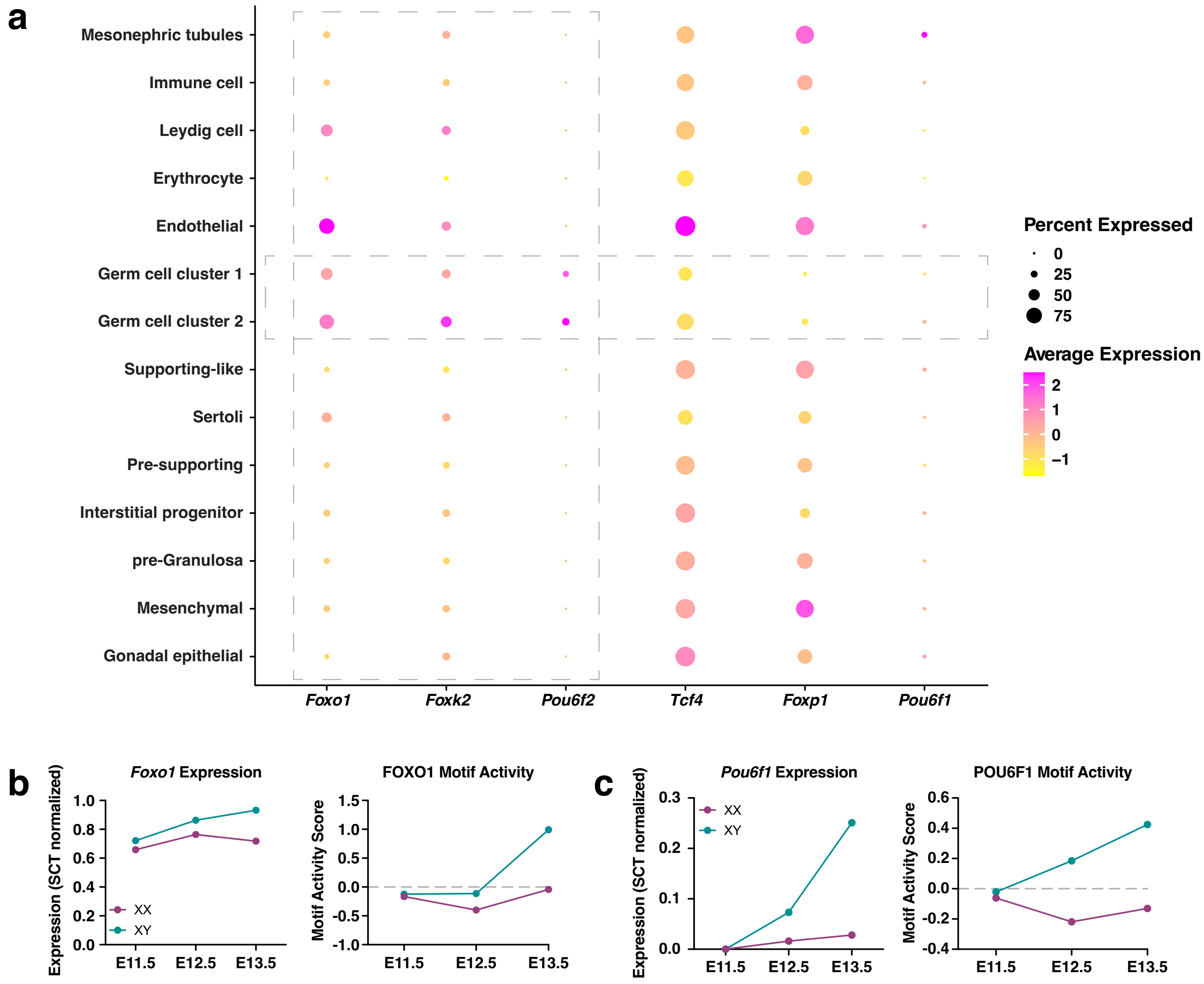


**Supplementary Figure 11: Identification of compelling transcription factor candidates enriched in XY PGCs.** (a) a) Dotplot of the average expression and percentage of cells expressing XY PGC-enriched transcription factors (TFs) in PGCs and gonadal somatic cells. Dashed boxes indicate TFs that are enriched in PGCs when compared to the somatic compartment. The color scale represents the average expression level, and the size of the dot represents the percentage of cells expressing the gene. (b-c) Line plots showing the expression levels and *in silico* chromatin binding, termed 'motif activity', of *Foxo1* (b) and *Pou6f1* (c) in E11.5-E13.5 XX (pink) and XY (teal) PGCs.

**Supplementary Figure 12**

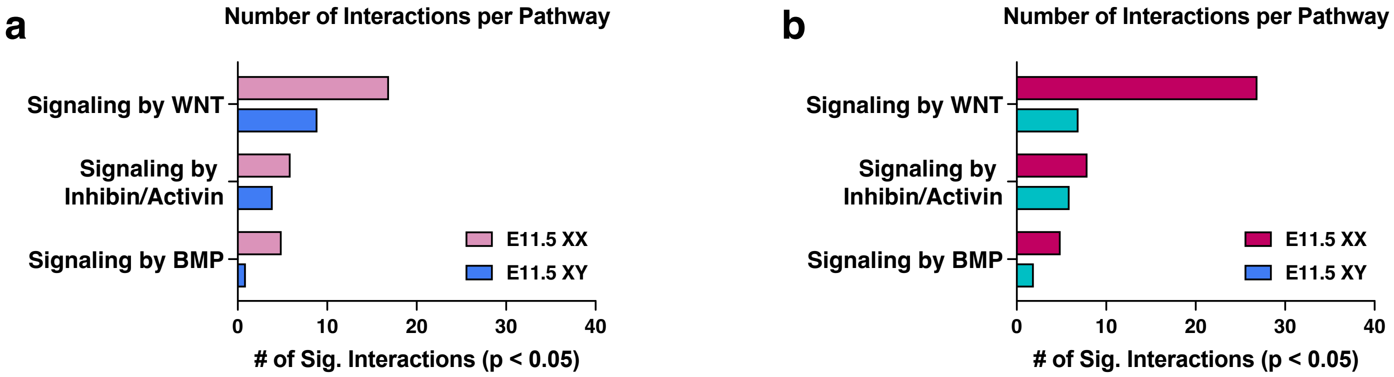


**Supplementary Figure 12: Number of interactions per signaling by WNT, inhibin/activin, and BMP pathways in XX and XY gonads.** (a-b) Number of significantly enriched ligand-receptor pairs, or interactions, per the signaling by WNT, inhibin/activin, and BMP pathways in E11.5 (a) and E12.5 (b) XX and XY gonads.
